# Reprogramming the neuroblastoma epigenome with a mitochondrial uncoupler

**DOI:** 10.1101/2021.09.05.459035

**Authors:** Haowen Jiang, Rachel L Greathouse, Bo He, Yang Li, Albert M. Li, Balint Forgo, Michaela Yip, Allison Li, Moriah Shih, Selene Banuelos, Meng-Ning Zhou, Joshua J. Gruber, Hiroyuki Shimada, Bill Chiu, Jiangbin Ye

## Abstract

Dysregulated DNA methylation is associated with poor prognosis in cancer patients, promoting tumorigenesis and therapeutic resistance^1^. DNA methyltransferase inhibitors (DNMTi) reduce DNA methylation and promote cancer cell differentiation, with two DNMTi already approved for cancer treatment^2^. However, these drugs rely on cell division to dilute existing methylation, thus the ‘demethylation’ effects are achieved in a passive manner, limiting their application in slow-proliferating tumor cells. In this study we use a mitochondrial uncoupler, niclosamide ethanolamine (NEN), to actively achieve global DNA demethylation. NEN treatment promotes DNA demethylation by activating electron transport chain (ETC) to produce α-ketoglutarate (α-KG), a substrate for the DNA demethylase TET. In addition, NEN inhibits reductive carboxylation, a key metabolic pathway to support growth of cancer cells with defective mitochondria or under hypoxia. Importantly, NEN treatment reduces 2-hydroxyglutarate (2-HG) generation and blocks DNA hypermethylation under hypoxia. Together, these metabolic reprogramming effects of NEN actively alter the global DNA methylation landscape and promote neuroblastoma differentiation. These results not only support Warburg’s original hypothesis that inhibition of ETC causes cell de-differentiation and tumorigenesis, but also suggest that mitochondrial uncoupling is an effective metabolic and epigenetic intervention that remodels the tumor epigenome for better prognosis.

## Materials and methods

### Cell lines and culture

CHP134, SK-N-BE(2) and NB16 cells were obtained from Dr. John M. Maris’ laboratory (Children’s Hospital of Philadelphia). SMS-KCNR cells were obtained from Dr. C. Patrick Reynolds’ laboratory (Texas Tech University Health Sciences Center)^3^. Certificate of analysis is available from each group. Ovarian cancer cell line OVCAR3 was from Dr. Erinn Rankin’s laboratory (Stanford University), and lung cancer cell lines H29 and H82 was from Dr. Julien Sage’s laboratory (Stanford University). All cell lines were cultured in DMEM/F12 medium (Caisson Labs, DFL15) supplemented with 1% Penicillin-Streptomycin (Gibco, 15140122), 10% FBS (Sigma, F0926) and extra 1mM glutamine (Gibco, 25030081).

### Protein extraction and immunoblot

Cells were washed with ice-cold PBS buffer (Invitrogen, 20012050) and lysed with harvest buffer (10mM Hepes pH 7.9, 50mM NaCl, 500mM sucrose, 0.1mM EDTA and 0.5% Trition X-100) supplemented with 1% Halt inhibitors (Thermo Scientific™, 78443) for 10 minutes. Cell lysate was centrifuged at 3000 rpm for 3 min at 4 °C. The supernatant containing cytosolic proteins was transferred to a different tube. The pellet containing nuclear proteins was dissolved with nuclear lysis buffer (10mM Hepes pH 7.9, 0.5M NaCl, 0.1mM EDTA, 0.1mM EGTA and 0.1% NP40) supplemented with 1% Halt inhibitors (Thermo Scientific™, 78443) for 5 minutes at 4 °C. Then the solution was sonicated at 4 °C for 6 cycles (1cycle = 30 s sonication and 30 s cooldown with “High” energy intensity) using a Bioruptor® Plus sonication (Diagenode, UCD-300). After sonication, the solution was centrifuged at 15,000 rpm for 10 minutes. The supernatant was kept as nuclear extraction. Protein concentration was determined by BCA assay (Thermo Scientific, 23227). 5 μg nuclear proteins or 7.5 μg cytosolic proteins were boiled in loading buffer with reducing reagents (Invitrogen, NuPAGE™ LDS Sample Buffer (4X), NP0007), then separated with SDS-PAGE (Nupage™ 4-12% Bis-Tris Protein Gels, 1.0 mm, Invitrogen™, NP0322BOX). Protein was transferred onto a nitrocellulose membrane (Thermo Scientific™, 88018). After blocking in 5% non-fat milk for 30 mintutes at room temperature (RT), the primary antibody was added for incubation overnight at 4 °C. After washing with TBST (CST, #9997, 1:10 diluted by deionized water) for 3 times, HRP-conjugated secondary antibodies were applied. Signals were detected with an ECL kit (Thermo Scientific™, 34577) by using BioRad Universal Hood II.

### RNA isolation, reverse transcription, and real-time PCR

The procedure was performed as described before^4^. Briefly, total RNA was isolated from 60mm tissue culture plates according to the TRIzol Reagent (Invitrogen, 15596026) protocol. 3 μg of total RNA was used in the reverse transcription reaction using the iScript cDNA synthesis kit (Bio-Rad). Quantitative PCR amplification was performed on the Prism 7900 Sequence Detection System (Applied Biosystems) using Taqman Gene Expression Assays (Applied Biosystems). Gene expression level were normalized to 18S rRNA.

### Cell proliferation and clonogenic assay

2 × 10^4^ cells were seeded in 12 well-plate and allowed to attached overnight. Then the cells were treated with various conditions as indicated for 48 h or 96 h (n = 3). After treatment, cells were counted using hemocytometer.

For clonogenic assay, 500 cells were plated in 60mm dishes. After 24h, cells were treated with indicated conditions. After 14 days, the cells were fixed by 50% methanol in PBS for 15 minutes and stained by crystal violet staining solution (0.5% crystal violet and 25% methanol in PBS). The quantification was done by using countPHICS ^5^.

### Neurite outgrowth assay

The quantification of neurite was performed as described before^6^. Shortly, 1 × 10^4^ SK-N-BE(2) and NB16 cells were plated in a 12 well-plate per well. After overnight incubation, cells were treated with various conditions as indicated for 72h. Then, images were taken by a Leica Florescent Microscope DMi8 in phase contrast mode (20× magnification). The lengths of the neurites were traced and quantified using the ImageJ plugin NeuronJ^7^. For each sample, total neurite length was measured and normalized to the number of cell bodies, mean value from 3 biological replicates was reported.

### Immunofluorescence staining

SK-N-BE(2) and NB16 cells were seeded into 8-champer slides (Thermo Fisher Scientific, 154534) with a density of 6×10^3^ cells/well overnight and treated with indicated treatment for 72h. Cells were fixed with 4% PFA in PBS with 0.1% Tween-20 at RT for 30 minutes followed by permeabilization with 0.1%Triton X-100 in PBS at room temperature for 10 min. The cells were washed with PBS twice and blocked with 2.5% horse serum in PBS at RT for 1 h. Then, cells were subjected to immunofluorescence staining with primary antibody at 4 °C overnight. After two washes with PBS, cells were incubated with Alexa Fluor 594-conjugated anti-rabbit secondary antibody (Life Technologies) at RT for 1 h followed by staining with DAPI (Vector Laboratories, H-1800-2). Images were acquired with Leica DMi8 microscope.

### RNA Sequencing and bioinformatics analysis

The total RNA from treatment groups (Ctrl and NEN treated, n = 3) were extracted using Trizol reagent according to the manufacturer’s instructions. The RNA-seq library was constructed and subjected to 150 bp paired-end sequencing on an Illumina sequencing platform (Novogene). RNA-seq analysis was performed using the kallisto and sleuth analytical pipeline^8-9^. In brief, a transcript index was generated with reference to Ensembl version 67 for hg19. Paired-end mRNA-seq reads were pseudo-aligned using kallisto (v0.42.4) with respect to this transcript index using 100 bootstraps (-b 100) to estimate the variance of estimated transcript abundances. Transcript-level estimates were aggregated to transcripts per million (TPM) estimates for each gene, with gene names assigned from Ensembl using biomaRt. Differential gene expression analysis was performed using the sleuth R package across pairwise groups using Wald tests, with significant hits called with a sleuth q-value < 0.05 and log2(fold change) > 0.693 or <-0.693. Gene set enrichment analysis (GSEA)^10-11^ was used to examine the significantly enriched pathways by comparing the normalized data of the entire RNA seq TPM dataset between groups as indicated to Molecular Signatures Database (MSigDB v7.4). All the annotated transcripts (∼21,427 features in total) with expression values were uploaded to a locally-installed GSEA tool (version 4.1.0) and compared against the H: hallmark gene sets (50 gene sets).

To generate the favorable or unfavorable prognosis gene sets, Kaplan-Meier assay were employed for all the genes independently by using all 11 available neuroblastoma patient databases from R2 (https://hgserver1.amc.nl/cgi-bin/r2/main.cgi). Each gene that passes filter of Kaplan-Meier assay showed significant correlation with prognosis (p-value<0.05) in relevant databases.

For functional annotation, differentially expressed genes were submitted to online Database for Annotation, Visualization and Integrated Discovery (DAVID) website (https://david.ncifcrf.gov/home.jsp) ^12-13^. The Analyses in DAVID were performed using the default parameters.

### Global DNA methylation analysis

For dot blot, 3 × 10^5^ cells were plated in 60mm dishes, after overnight incubation, the cells were treated with DMSO, 1µM NEN or 3.5mM dimethyl α-ketoglutarate (DMKG) for 1h or 3h. Cells were scaped in ice-cold PBS, the cell pellet were collected by centrifuging at 3000rpm for 3 minutes. DNA samples were extracted by using PureLink™ Genomic DNA Mini Kit (Gibco, K182001), then, they were denatured at 95 °C for 10 minutes and placed on ice immediately for at least 5 minutes before multiple dilutions. Samples at different dilutions were immobilized on a nitrocellulose membrane (Thermo Scientific™, 88018). Subsequently, the membranes were air-dried for 20minutes, UV cross-linked (UV stratalinker 1800, auto-crosslink mode), blocked with 5% non-fat milk for 1h at RT followed by 16h incubation at 4 °C with anti-5-methylcytosine (5mC) antibodies. After three washes in TBST, HRP-conjugated secondary antibodies were added and incubated for 1h at RT. Signals were detected using an ECL kit (Thermo Scientific™, 34577) on BioRad Universal Hood II. The membranes were then stained with methylene blue (0.02% methylene blue in 300Mm Sodium acetate, Ph 5.3) to confirm equal DNA loading.

In addition, 5 × 10^5^ SK-N-BE(2) cells were plated in 60mm dishes overnight, then the cells were treated by DMSO,1µM NEN or 5-AZA for 3h, 6h, 24h and 48h, respectively. DNA from the cells were collected and performed DNA extraction as described above. Quantitative analysis of global 5-mC levels was carried out with the MethylFlash Methylated DNA Quantification Kit (Epigentek, P-1030-48) according to the manufacturer’s instructions. Results are presented as the percentage of methylated DNA (5-mC) to total DNA.

### Stable isotope tracing analysis

For glutamine isotope tracing under normoxia, SK-N-BE(2) and NB16 cells were pretreated with DMSO or 1 μM NEN for 3h. Then the culture medium was changed to DMEM/F-12, no glutamine (Gibco, 21331020) with 4mM U-^13^C_5_-glutamine (Cambridge Isotope Laboratories, CLM-1822-H) and 10% dialyzed FBS (Gibco, 26400044) under same treatment condition as above for 2h. For glutamine isotope tracing under hypoxia, the SK-N-BE(2) cells were pretreated by DMSO or 1 μM NEN under normoxia or hypoxia for 4h. Then changed the medium to DMEM/F-12, no glutamine (Gibco, 21331020) with 4mM 1-^13^C-glutamine (Cambridge Isotope Laboratories, CLM-CLM-3612-PK) and 10% dialyzed FBS (Gibco, 26400044) for 2h with same treatment under normoxia or hypoxia for 2h.

### Liquid chromatography-mass spectrometry analysis

For the metabolomics and isotope tracing analysis, cells were washed with cold PBS, lysed in 80% Ultra LC-MS acetonitrile (Fisher Scientific, A955-4) on ice for 15 minutes, centrifuged at 20,000 g for 10 minutes at 4 °C, and the supernatant was subjected to mass spectrometry analysis. Liquid chromatography was performed using an 1290 Infinity LC system (Agilent, Santa Clara, US) coupled to a Q-TOF 6545 mass spectrometer (Agilent, Santa Clara, US). A hydrophilic interaction chromatography method (HILIC) with a BEH amide column (100 × 2.1 mm i.d., 1.7 μm; Waters) was used for compound separation at 35 °C with a flow rate of 0.3 ml/ min. The mobile phase A consisted of 25 mM ammonium acetate and 25 mM ammonium hydroxide in water and the mobile phase B was acetonitrile. The gradient elution was 0–1 min, 85% B; 1–12 min, 85% B → 65 % B; 12–12.2 min, 65 % B → 40%B; 12.2–15 min, 40%B. After the gradient, the column was re-equilibrated at 85%B for 5 min. The overall runtime was 20 min, and the injection volume was 5 μL. Agilent Q-TOF was operated in negative mode and the relevant parameters were as follows: ion spray voltage, 3500 V; nozzle voltage, 1000 V; fragmentor voltage, 125 V; drying gas flow, 11 L/min; capillary temperature, 325 °C, drying gas temperature, 350 °C; and nebulizer pressure, 40 psi. A full scan range was set at 50 to 1600 (m/z). The reference mass was 119.0363 and 980.0164. The acquisition rate was 2 spectra/s. Targeted analysis, isotopologues extraction and natural isotope abundance correction were performed by t Profinder B.10.00 (Agilent, Santa Clara, USA).

### Mouse orthotopic neuroblastoma model

All mouse procedures were performed in accordance with Stanford University recommendations for the care and use of animals and were maintained and handled under protocols approved by the Institutional Animal Care and Use Committee. All procedures were performed with female NC nude mice (Taconic, Hudson, NY, USA) at 7 weeks of age. Procedures and ultrasound measurements (see below) were performed under general anesthesia using isoflurane inhalation. Orthotopic tumors within the adrenal gland were created as described before^14^. Briefly, a transverse incision was made on the left flank to locate the left adrenal gland, and 2 mL of phosphate buffered saline (PBS) containing 10^4^ SK-N-BE(2) cells were injected into the adrenal gland using a 30G needle. Fascia and skin were closed in separate layers. Tumor formation was monitored by non-invasive ultrasound measurements, and the animals euthanized when the tumor volume exceeded 1,000 mm^3^.

### Monitoring tumor growth with high frequency ultrasound

After securing the mouse in a prone position, a Visual Sonics Vevo 2100 sonographic probe (Toronto, Ontario, Canada) was applied to the left flank to locate the left adrenal gland and the tumor. Serial cross-sectional images (0.076 mm between images) were taken. The tumor volume was measured using the 3-D reconstruction tool (Vevo Software v1.6.0, Toronto, Ontario, Canada).

### Immunohistochemistry

Unstained sections from the study cases were heated for 30 minutes in Bond™ Epitope Retrieval Solution 2 (No. AR9640; Leica Biosystems) using Leica BOND-MAXTM (Leica Microsystems), and incubated with either anti-MYCN mouse monoclonal antibody (NCM II 100at a dilution of 1:200) or anti-human MYC rabbit monoclonal antibody(clone Y69; #1472–1; Epitomics at a dilution of 1:200) in Bond™ Primary Antibody Diluent (No. AR9352; Vision BioSystems). The counter staining with hematoxylin was performed for MYC protein staining slides, but no counterstaining was performed for MYCN protein staining slides.

### Statistics

For cell proliferation and MS experiments, three biological repeats were used for data analysis. Results were represented as mean ± SEM or mean ± SD. The student’s t-test was performed to determine the significance between groups (two-tailed, unequal variance).

## Introduction

Growing evidence points to the critical role of dysregulated epigenetics in promoting cancer progression^15-18^. Particularly, CpG islands hypermethylation in promoter region leads to the silencing of tumor suppressors, cell differentiation markers associated with poor prognosis in cancer patients, and therapeutic resistance ^19-22^.

Unlike genetic mutations, epigenetic changes are reversible. Thus, CpG island hypermethylation has become an attractive target for cancer therapy. The DNA methyltransferase inhibitors (DNMTi) azacitidine and decitabine have shown promising effect in clinical trials and have been approved by the FDA and EMA to treat hematopoietic malignancies ^23-25^. Nonetheless, there are several limitations of using DNMTi in clinic. First, they only block methylation on newly synthesized DNA strands rather than remove methyl groups from existing methylated CpG islands^26^. Second, DNMTi are considered as week mutagen since they incorporate into DNA ^26^. Third, they cause myelosuppression and gastrointestinal toxicities ^2^. These limitations prompt a need to identify more effective and safer strategies to target DNA hypermethylation for cancer therapy.

DNA demethylation in mammals is achieved through TET enzyme-mediated sequential oxidation of 5-methylcytosine (5-mC) to 5-hydroxymethylcytosine (5-hmC), 5-formylcytosine (5-fC) and then 5-carboxylcytosine (5-caC), followed by thymine DNA glycosylase (TDG)-mediated excision of 5-fC and 5-caC coupled with base excision repair^27^. This multi-step DNA demethylation process requires the co-substrate α-ketoglutarate (α-KG), a tricarboxylic acid (TCA) cycle intermediate that can be generated from glucose or glutamine^28^. However, under hypoxia, α-KG is reduced and converted to 2-hydroxyglutarate (2-HG) as a an oncometabolite due to a reduced NAD^+^/NADH ratio ^29-33^, thus inhibiting TET ^34^. Based on these discoveries, we hypothesize that DNA hypermethylation would be reversed by increasing the intracellular NAD^+^/NADH ratio, thus, restoring the redox homoeostasis. Consequently, the aberrant metabolic and epigenetic phenotypes of cancer cells could be corrected.

The electron transport chain (ETC) is the major site for cells to regenerate NAD^+^ from NADH. We hypothesize that ETC activation by mitochondrial uncoupler would increases the NAD^+^/NADH ratio, leading to an increase of the αKG/2HG ratio. One of them, niclosamide ethanolamine (NEN) facilitates proton influx across the mitochondrial inner membrane without generating ATP to activate the electron transport chain (ETC) ^35-36^. In this study, we use NEN to reprogram the metabolism and epigenetic landscape of cancer cells. We have found that NEN treatment increases the intracellular NAD^+^/NADH ratio, αKG/2HG ratio, and pyruvate/lactate ratio. This metabolic reprogramming reduces global DNA methylation and activates differentiation-related gene expression. N-Myc, an oncogenic transcription factor often amplified in neuroblastoma (NB) and is associated with worse patient survival, is downregulated by NEN treatment. Furthermore, NEN treatment inhibits reductive carboxylation, a key metabolic pathway that converts α-KG to citrate to support cancer cell growth upon ETC inhibition^31, 37^. Importantly, NEN treatment reduces 2HG-mediated DNA hypermethylation under hypoxia. Together, these results uncover a new role of mitochondrial uncoupling as an epigenetic intervention that promotes DNA demethylation and tumor cells differentiation.

## Results

### Mitochondrial uncoupling increased intracellular α-KG and α-KG/2-HG ratio

Mitochondrial uncoupling is a process that dissipates the proton gradient across the inner mitochondrial membrane, activating the ETC to promote NADH oxidation^38-40^. Niclosamide ethanolamine (NEN) is a salt form of the FDA-approved mitochondrial uncoupler drug niclosamide with an excellent safety profile^41-44^. As expected, NEN treatment increased ADP/ATP and AMP/ATP ratios (Figure S1a, b). Importantly, NEN treatment increased intracellular NAD^+^/NADH ratio in both SK-N-BE(2) and NB16 cells (Figure 1a, b). In addition, the pyruvate/lactate ratio, determined by the NAD^+^/NADH ratio^45^, was also increased upon NEN treatment, indicating inhibition of the Warburg effect (Figure 1a, b). Furthermore, NEN treatment did not cause oxidative stress in our experimental setting based on the observation that NEN did not reduce the glutathione (GSH)/glutathione disulfide (GSSG) ratio in neither SK-N-BE(2) nor NB16 cells (Figure S1a, b).

**Figure 1.**
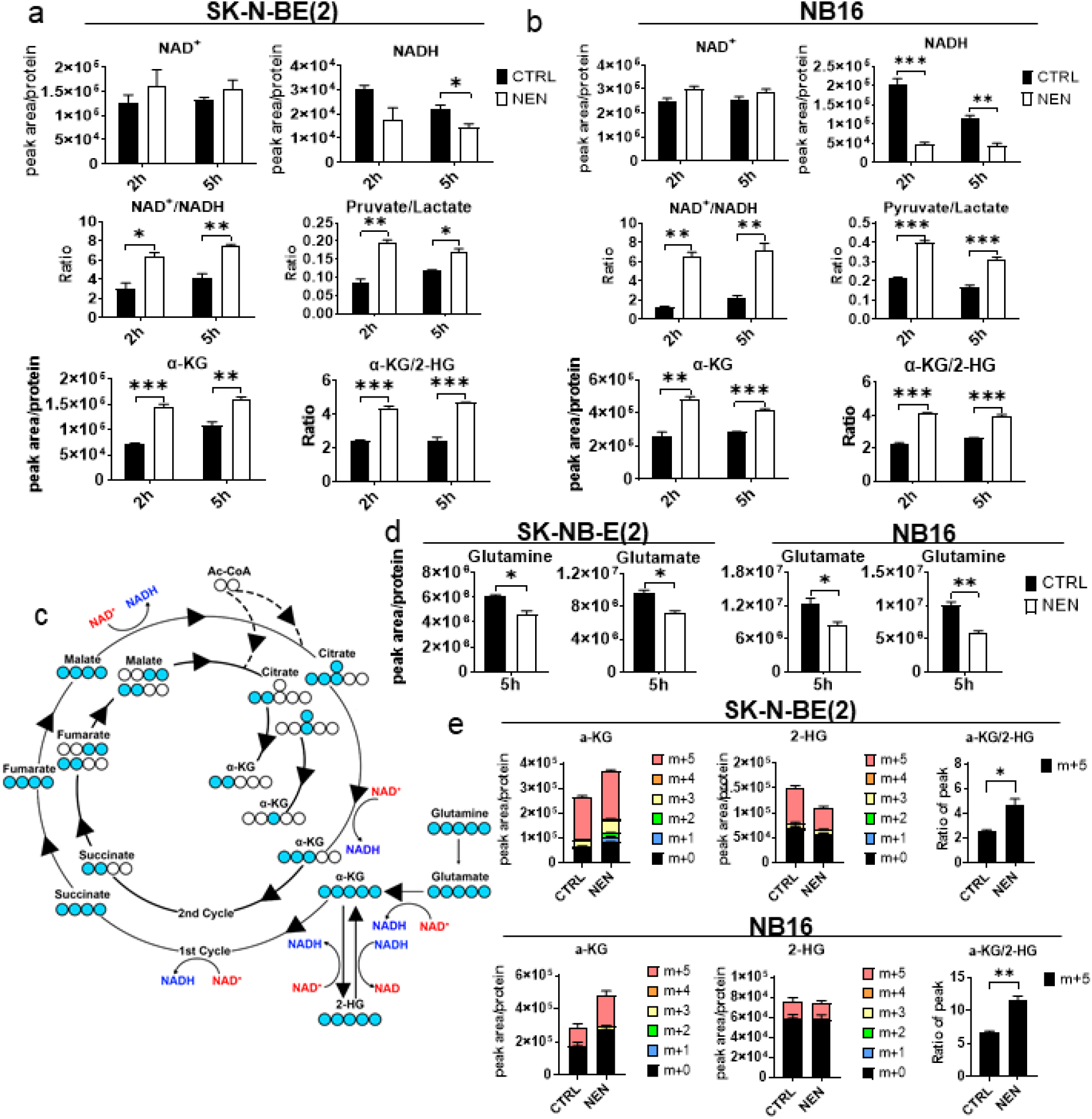
NEN treatment accelerates glutaminolysis to increase cellular αKG. Relative intracellular metabolite levels were measured using LC/MS in SK-N-BE(2) cells (**a**) and NB16 cells (**b**) treated with DMSO or 1 μM NEN for 2h or 5h. (**c**) Relative intracellular L-glutamine and L-glutamate levels were measured using LC/MS in SK-N-BE(2) cells and NB16 cells treated with DMSO or 1 μM NEN for 5h. **(d)** Schematic of ^13^C-labeling patterns of TCA cycle metabolites froi U-^13^C-glutamine. **(e)** SK-N-BE(2) and NB16 cells were pretreated by DMSO or 1 μM NEN for 3h, then labeled with U-^13^C-glutamine for 2h. Relative isotopic labelling abundance in αKG, 2HG and the ratio of m+5 αKG/m+5 2HG were measured using LC/MS. Data represent mean ± SEM (n = 3, biologically repeats). Representative of at least two independent experiments. *P < 0.05, **P < 0.01, ***P < 0.001. Two-sided Student’s t-test.

Because multiple key reactions in TCA cycle use NAD^+^ as an electron acceptor, a high NAD^+^/NADH ratio is the major driving force for TCA cycle (Figure 1c). We next examined how mitochondrial uncoupling regulates TCA cycle metabolite levels. As the product of the first step of TCA cycle, citrate was not affected by NEN treatment (Figure S1). Surprisingly, intracellular α-KG levels increased significantly in both cell lines after NEN treatment (Figure 1a, b). Succinate also accumulated under NEN treatment while the αKG/succinate ratio stayed unchanged (Figure S1a, b), and no significant increase of either fumarate or malate was observed after NEN treatment (Figure S1a, b). Aspartate, which is derived from oxaloacetate, also accumulated after NEN treatment (Figure S1a, b). Because glutamine and glutamate provide the carbon backbone to generate α-KG, we wondered whether the increased α-KG originated from glutamine. NEN treatment significantly reduced the intracellular glutamine and glutamate levels, suggesting the acceleration of glutaminolysis by mitochondrial uncoupling (Figure 1d).

To determine how mitochondrial uncoupling alters the glutamine flux into TCA cycle, we carried out [U-^13^C_5_]-glutamine tracing assay (Figure 1c). Surprisingly, in SK-N-BE(2) cells, no significant increase of m+5 α-KG was observed. However, the labelling percentage of m+3 (2^nd^ cycle) and m+1/2(3^rd^ cycle) α-KG significantly increased upon NEN treatment (Figure 1d,e). Also, the labelling abundance of m+2(2^nd^ cycle) and m+1/2(3^rd^ cycle) succinate and aspartate (from oxaloacetate) significantly increased under NEN treatment (Figure S2a), suggesting that NEN treatment accelerates the of TCA cycle flux in the oxidative direction. Importantly, the m+5 labelling abundance of 2-HG was significantly decreased by NEN treatment in SK-N-BE(2) cells (Figure 1e). The ratio of m+5 α-KG/m+5 2-HG decreased under NEN treatment (Figure 1e), indicating that NEN inhibits the conversion of α-KG to 2-HG. In NB16 cells, no significant reduction of m+5 2-HG was observed, possibly due to the lower 2-HG generation in this cell line (Figure 1e). The increased m+5 and m+3 labeling from glutamine accounted for the α-KG increase (Figure 1e), and was associated with increased m+4 succinate and m+4 aspartate (Figure S2b). We also tested the metabolic reprograming effect of NEN on other cancer cell lines, including an ovarian cancer cell line OVCAR3, and two lung cancer cell lines H29 and H82. All the cell lines showed similar metabolic reprograming effects, featured by the increased ADP/ATP ratio and α-KG/2-HG ratio, indicating that this is a universal metabolic reprogramming effect shared by multiple cancer types (Figure S3). Together, these data suggest that NEN treatment upregulates cellular α-KG through two potential mechanisms: accelerating glutaminolysis and blocking the conversion of α-KG to 2-HG.

### Mitochondrial uncoupling decreases global DNA methylation and promotes neuroblastoma cell differentiation

Increased intracellular α-KG levels due to mitochondrial uncoupling may promote DNA demethylation. Indeed, NEN treatment reduced global DNA methylation in NB16 cells, as determined by dot blot (Figure 2a). To directly increase intracellular α-KG, a cell-permeable form of α-KG, dimethyl α-ketoglutarate (DMKG) were used, and it also reduced global DNA methylation (Figure 2a). To capture the dynamic changes of global DNA methylation, we used an ELISA-based kit to quantify the DNA methylation in cells treated by NEN or a DNMTi 5-Azacytidine (5-Aza) at various time points. We found that NEN reduced the global DNA methylation faster at early time points (1h and 3h) than 5-Aza (Figure 2b), possibly because NEN promotes active demethylation instead of passively blocking methylation, a process that requires cell division and DNA replication.

**Figure 2.**
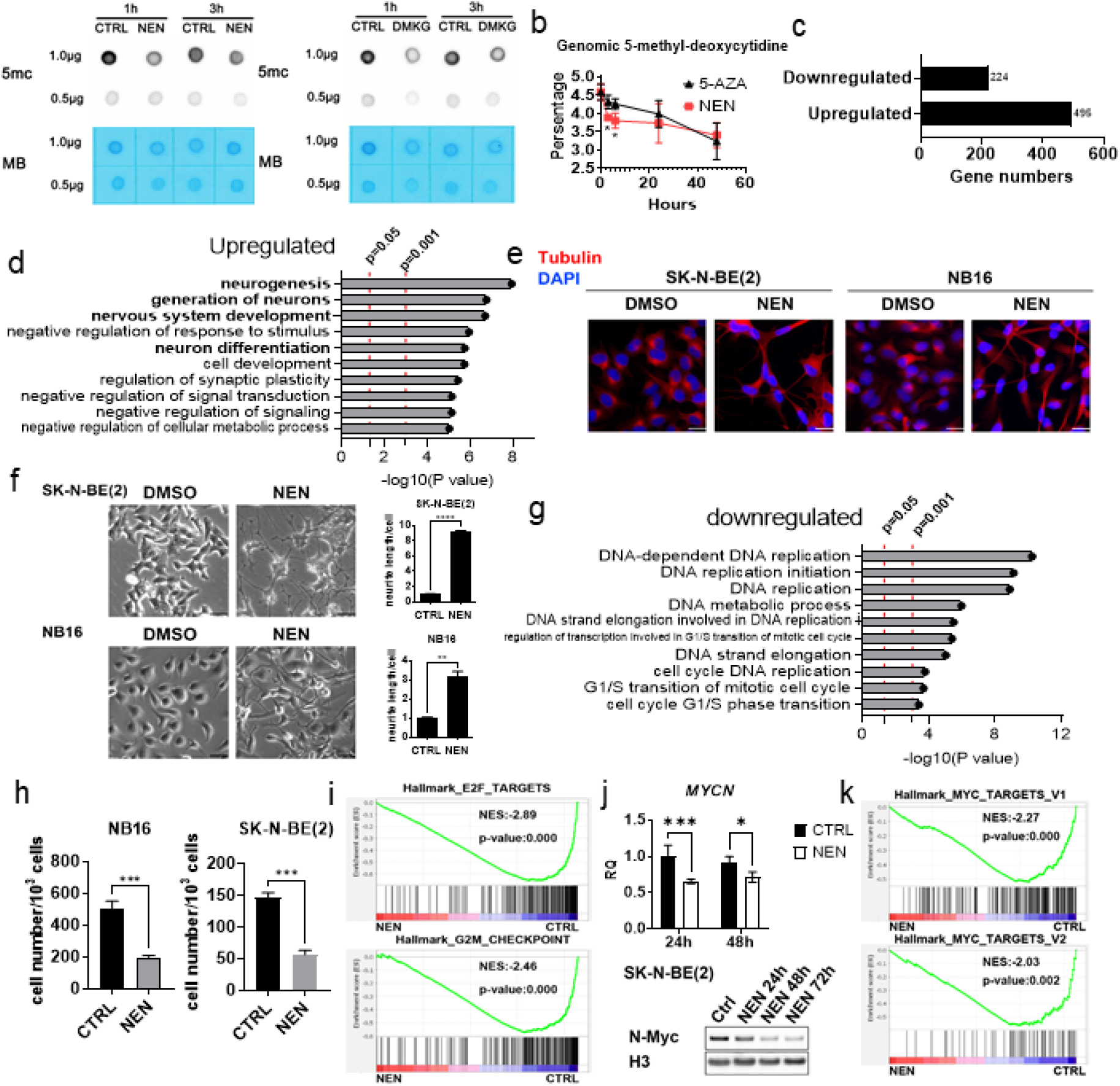
NEN decreased global DNA methylation and promote neuron differentiation. (a) Global DNA methylation in NB16 cells treated with DMSO, 1 μM NEN or 3.5mM DMKG for 1hrs and 3hrs was measured using dot blot. (b) Quantification of global DNA methylation in SK-N-BE(2) cells treated by DMSO, 1 μM NEN or 1 μM 5-AZA for 3h, 6h, 24h and 48h by using the MethylFlash™ Global DNA Methylation (5-mC) ELISA Easy Kit. (c) Number of genes differentially expressed after NEN treatment for 16hrs in SK-N-BE(2) cells with a sleuth q-value < 0.05 and fold change estimate b > abs(ln(2)). (d) The top 10 gene expression signature pathways enriched from upregulated genes from David analysis. (e) Immunofluorescence staining of β-tubulin III(Red) and DAPI (Blue) in SK-N-EB(2) and NB16 cells treated by DMSO, 1 μM NEN for 72h. Scale bar: 25μm. (f) Left: morphological feature of NB16 and SK-N-BE(2) cells treated by DMSO or 1 μM NEN for 96h (Scale bar: 50 μM). Right: Quantification of neurite outgrowth with NeuronJ. (g) The top 10 gene expression signatures enriched from downregulated genes from David analysis. (f) Cells were plated in 12 wells plates (2×10^4^ cells/well). After 24hrs, cells were treated with DMSO or 1 μM NEN for 3 days, and then counted. (i) GSEA of E2F targets and G2M checkpoints pathways genes from the RNA-seq (n=3) experiments in SK-N-BE(2) cells. (j) mRNA and protein levels of N-myc were examined in SK-N-BE(2) cells treated with 1μM NEN for indicated time. (k) GSEA of N-myc targets pathway genes from the RNA-seq (n=3) experiments in SK-N-BE(2) cells.

To investigate how NEN treatment alters the global gene expression profile, we performed RNA-seq in SK-N-BE(2) cells treated with DMSO or NEN for 16h. Corresponding to the DNA demethylation effect of NEN, 495 genes were significantly upregulated while 224 genes were downregulated by NEN treatment with a sleuth q-value < 0.05 and fold change estimate b > abs(ln(2)) (Figure 2c). The upregulated genes were enriched in multiple pathways including neurogenesis, nervous system development and neuron differentiation (Figure 2d). Consistent with the pathway enrichment result, NEN treatment induced neuron differentiation morphology change in both SK-NE-BE(2) and NB16 cells, as determined by the neurite length measurement and immunofluorescence staining against neuron differentiation marker β-tubulin III (Figure 2e and f). In contrast, the NEN-downregulated genes were enriched in pathways involved in DNA replication and cell cycle progression (Figure 2g). As expected, NEN treatment significantly inhibited cell proliferation (Figure 2h). In addition, Gene Set Enrichment Analysis (GSEA) of the RNA-seq results showed that NEN treatment significantly deviated from two important cell division hallmark “E2F-TARGETS” and “G2M_CHECKPOINT” when compared to control (CTRL) treatment (Figure 2i), indicating downregulation of cell division-related genes. Intriguingly, it was reported that E2Fs transcriptionally upregulates *MYCN* ^46^, the key oncogenic factor that is amplified in neuroblastoma and associated with poor patient outcome. NEN treatment significantly reduced the mRNA and protein level of N-Myc (Figure 3j). Consistent with the reduced N-Myc protein levels, GESA analysis showed that NEN treatment significantly deviated from hallmark “MYC_TARGETS_V1” and “MYC_TARGETS_V2” when compared to control (CTRL) treatment (Figure 3k), indicating that NEN reduced the expression N-Myc targeted genes.

**Figure 3.**
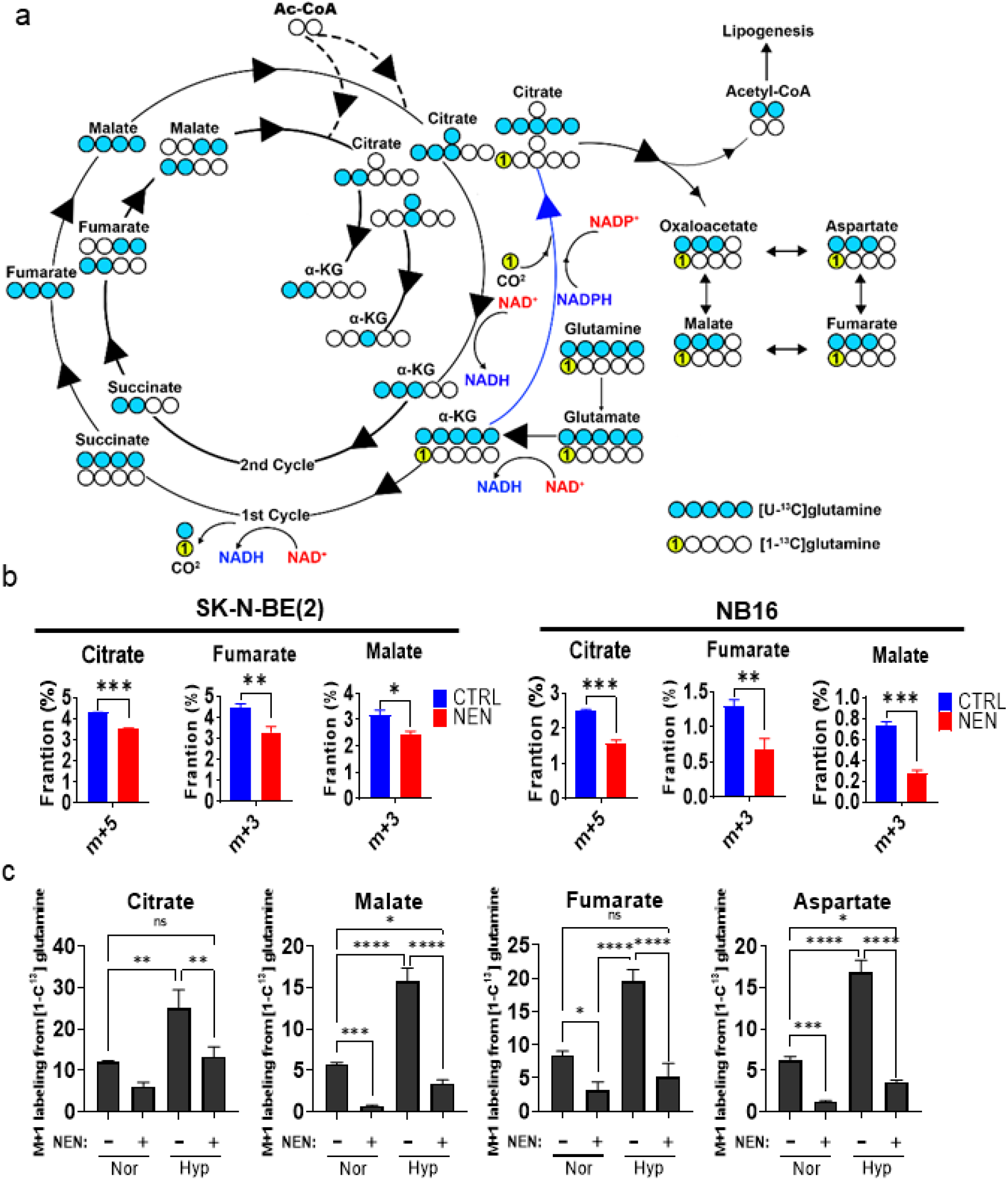
NEN treatment decreased the reductive carboxylation. (a) Schematic of carbon atom (circles) transitions from U-^13^C-glutamine(blue) and 1-^13^C glutamine (yellow) into oxidation TCA cycle or reductive carboxylation. (b) SK-N-BE(2) and NB16 cells were pretreated by DMSO or 1 μM NEN for 3h, then switched to medium contained same treatment in the presence of U-^13^C-glutamine for 2h. The Isotopomer distribution of citrate, malate, fumarate and aspartate from U-^13^C-glutamine were measured using LC/MS. (c) SK-N-BE(2) cells were pretreated by DMSO or 1 μM NEN for 3h under normoxia or hypoxia (0.5% oxygen), then switched to medium contained same treatment in the presence of 1-^13^C-glutamine for 3h.The Isotopomer distribution of citrate, malate and fumarate were measured using LC/MS. Data represent the mean ± SEM from three biologically repeats. *P < 0.05, **P < 0.01 and ***P < 0.001. Two-sided Student’s t-test..

### Mitochondrial uncoupling inhibits reductive carboxylation

Cancer cells under hypoxia or carrying mutations that suppress mitochondrial function display reductive carboxylation^47-49^, a reaction that converts α-KG to citrate to provide acetyl-CoA for lipid synthesis (Figure 3a). In the U-^13^C_5_-glutamine tracing assay, we found that NEN treatment significantly inhibited reductive carboxylation flux, as determined by the decreased labelling faction of m+5 citrate, m+3 fumarate, and m+3 malate (Figure 3b). We next tested whether mitochondrial uncoupling could also reverse reductive carboxylation under hypoxia. When [1-^13^C]-glutamine is used as a tracer, labelling carbon will only be detected in metabolites from the reductive carboxylation pathway (m+1 citrate, m+1 fumarate, m+1 malate, and m+1 aspartate) (Figure 3a). NEN treatment not only reduced basal reductive carboxylation flux under normoxia, but also fully repressed hypoxia-induced reductive carboxylation flux (Figure 3c), indicating that the mitochondrial uncoupler NEN is an effective inhibitor of reductive carboxylation.

### Mitochondrial uncoupling inhibits 2-HG generation and DNA hypermethylation under hypoxia

Clinically, tumor hypoxia is a significant obstacle to treatment because hypoxic tumor cells are more resistant to radiation ^50-51^ and chemotherapy ^52-54^. Under hypoxia, due to reduced NAD^+^/NADH ratio, α-KG is reduced and converted to L-2-HG ^29-30^, which may repress α-KG-dependent dioxygenase, including TET DNA demethylase. We found that NEN treatment could partially restore NAD^+^/NADH and pyruvate/lactate ratios under hypoxia (Figure 4a). Importantly, NEN treatment partially restored α-KG and significantly reduced 2-HG levels under hypoxia (Figure 4a). Our previous study showed that in neuroblastoma cells, hypoxia treatment inhibited neuroblastoma cell differentiation, which could not be restore by α-KG supplementation ^6^, possibly because supplemented α-KG was converted to 2-HG ^29^. Intriguingly, DNA hypermethylation induced by hypoxia could be repressed by NEN but not αKG, suggests that NEN supplementation has an advantage in demethylating DNA over α-KG supplementation under hypoxia (Figure 4b), because NEN treatment reduces while α-KG increases 2-HG under hypoxia (Figure 4c). In addition, NEN supplementation can induce differentiation under hypoxia (Figure 4d). Importantly, we found that 4 days hypoxia pretreatment increased cell proliferation when the cells were replated under normoxia condition (with no treatment). However, when cells were pretreated under hypoxia with NEN, this gain of proliferation advantage from hypoxia pretreatment is blocked (Figure 4c).

**Figure 4.**
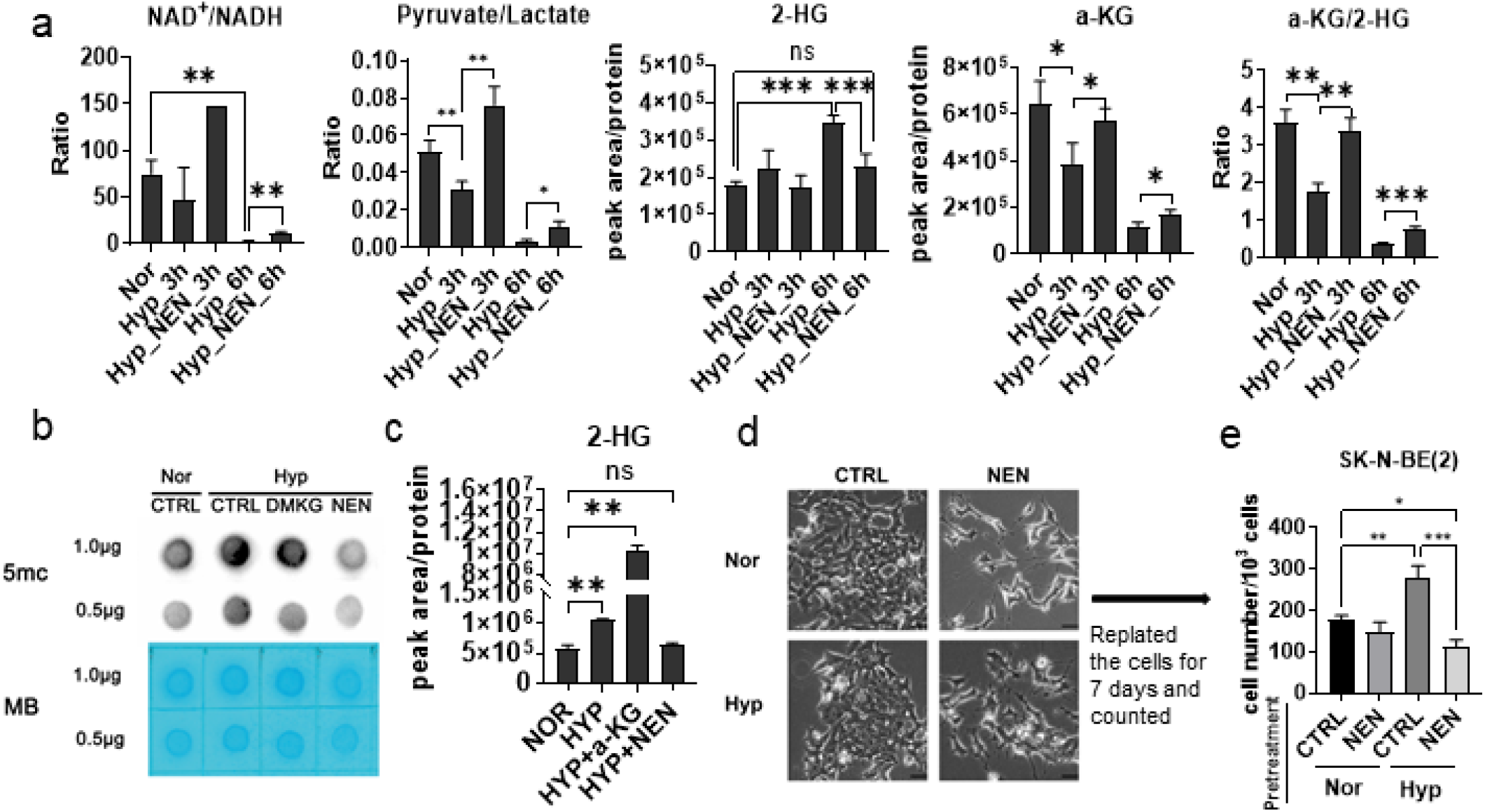
NEN inhibit 2-HG generation and DNA hypermethylation under hypoxia. (a) Relative Intracellular metabolite level or ratio were measured by LC/MS in SK-N-BE(2) cells treated by DMSO under normoxia or DMSO / 1 μM NEN under hypoxia for 3h or 6h. (b) Morphological feature of SK-N-BE(2) cells treated by DMSO or 1 μM NEN under normoxia or hypoxia for 4 days(Scale bar: 50 μM). (c) Trypsinized the cells from (b) and replated 2×104 cells/well were plated in 12 wells plates. Count the cells after 5 days. (d) Analysis and quantification of global DNA methylation in CHP134 cells treated by DMSO or 1 μM NEN or 3.5mM DMKG under normoxia or hypoxia for 6hrs by using dot blot. (e) Relative Intracellular 2-HG level were measured by LC/MS in CHP134 cells treated by DMSO or 1 μM NEN or 3.5mM DMKG under normoxia or hypoxia for 5hrs. Data are the mean ± SEM from n = 3 biologically independent. *P < 0.05 **P < 0.01 and ***P < 0.00 for comparisons were calculated using a two-sided Student’s t-test.

### NEN supplementation inhibits growth of orthotropic neuroblastoma *in vivo*

Next, we examined whether NEN has anti-growth effect *in vivo* using an orthotopic neuroblastoma xenograft model. Because intraperitoneal injection of NEN shows poor plasma pharmacokinetics (Figure S4a), we used diet which contain 2000ppm NEN for treatment delivery ^55^. LC/MS analysis indicated that dietary NEN supplementation led to abundant NEN accumulation in plasma (1-3µM) (Figure 5b). The NEN accumulation in xenograft tumors is comparable to levels found in the kidney, although lower than levels in the liver (Figure 5c). NEN supplementation significantly slowed tumor growth as compared to control group (Figure 5d). Furthermore, tumor cells from NEN-treated group had much fewer enlarged prominent nucleoli (Figure 5e), which indicates active ribosome biogenesis and worse prognosis in NB patients^56^. Importantly, N-Myc protein expression was significantly reduced in the NEN-treated xenografts (Figure 4f). This is consistent with our *in vitro* observation (Figure 2j and k).

**Figure 5.**
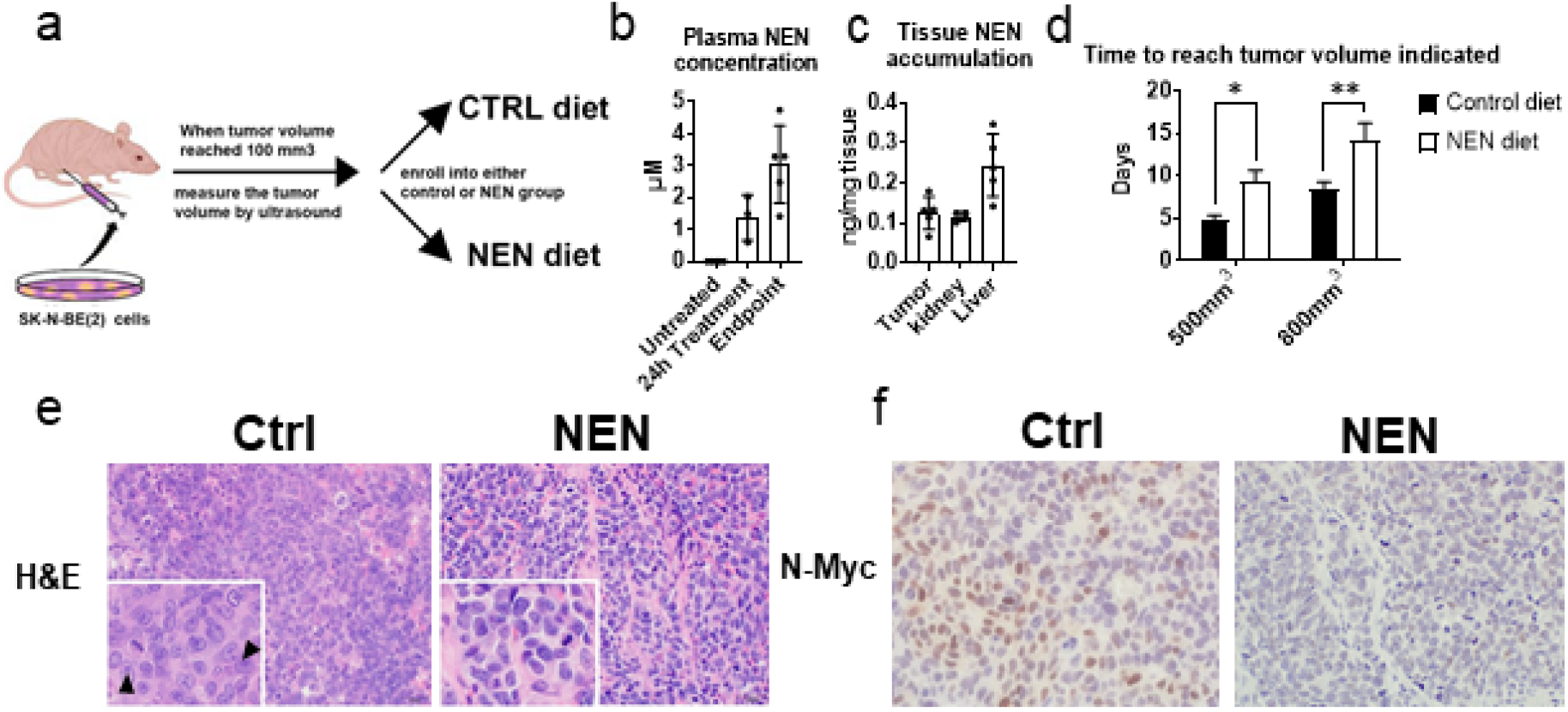
NEN supplementation inhibits growth of orthotropic neuroblastoma in vivo. (a) The schematic of in vivo experiment (b,c) The plasma and tissue NEN accumulation were measured by LC/MS. (d) Quantification of time to reach certain tumor volume on CTRL group(n=6) and NEN group (n=4). (e) Nucleolar formation in the tumors cells were examined by H&E staining in both CTRL and Treatment group. The arrows point out the prominent nucleolar formation. (f) The tumors in both CTRL and NEN treatment group were stained with H&E and processed for N-Myc immunohistochemistry staining.

### NEN-induced gene expression changes indicate favorable prognosis in neuroblastoma patients

To further explore the clinical relevance of NEN-induced gene expression profile changes, we integrated the gene expression signature of RNA-seq result from NEN treatment experiment and 11 neuroblastoma patient gene expression profile from R2 database. Using the datasets from R2 database, we generated gene sets that indicated favorable prognosis or unfavorable prognosis from the 11 neuroblastoma patient studies (The p-value of Kaplan-Meier analysis of each gene is less than 0.05). Consequently, these gene sets were defined as pathways for GSEA analysis. Surprisingly, NEN-upregulated genes are enriched in all the favorable prognosis gene sets except one (Figure 6a, b). In contrast, NEN-downregulated genes are enriched in unfavorable prognosis gene sets (Figure 6a, b). The only exception is that NEN-upregulated genes are enriched in an unfavorable prognosis gene set derived from N-Myc non-amplification neuroblastomas. A potential explanation for this discrepancy is that the RNA-seq data was generated from SK-N-BE(2) cells, which are N-Myc-amplified(Figure 2k). In addition, we constructed sets of overlapped genes from 7 gene sets that contain more than 1000 genes; namely, an overlapped “favorable prognosis gene set” with 286 genes and an overlapped “unfavorable prognosis gene set” with 270 genes (Figure 6c, d). These two genes set were submitted to pathway enrichment through online Database for Annotation, Visualization and Integrated Discovery (DAVID) analysis. The overlapped favorable prognosis gene sets were enriched for multiple neuron differentiation terms while the overlapped unfavorable prognosis gene sets were enriched for cell-cycle related terms (Figure 6c, d). Together these data suggest that NEN treatment reprograms the transcriptome to a gene expression profile that is associated with favorable prognosis in patients with N-Myc-amplified neuroblastoma.

**Figure 6.**
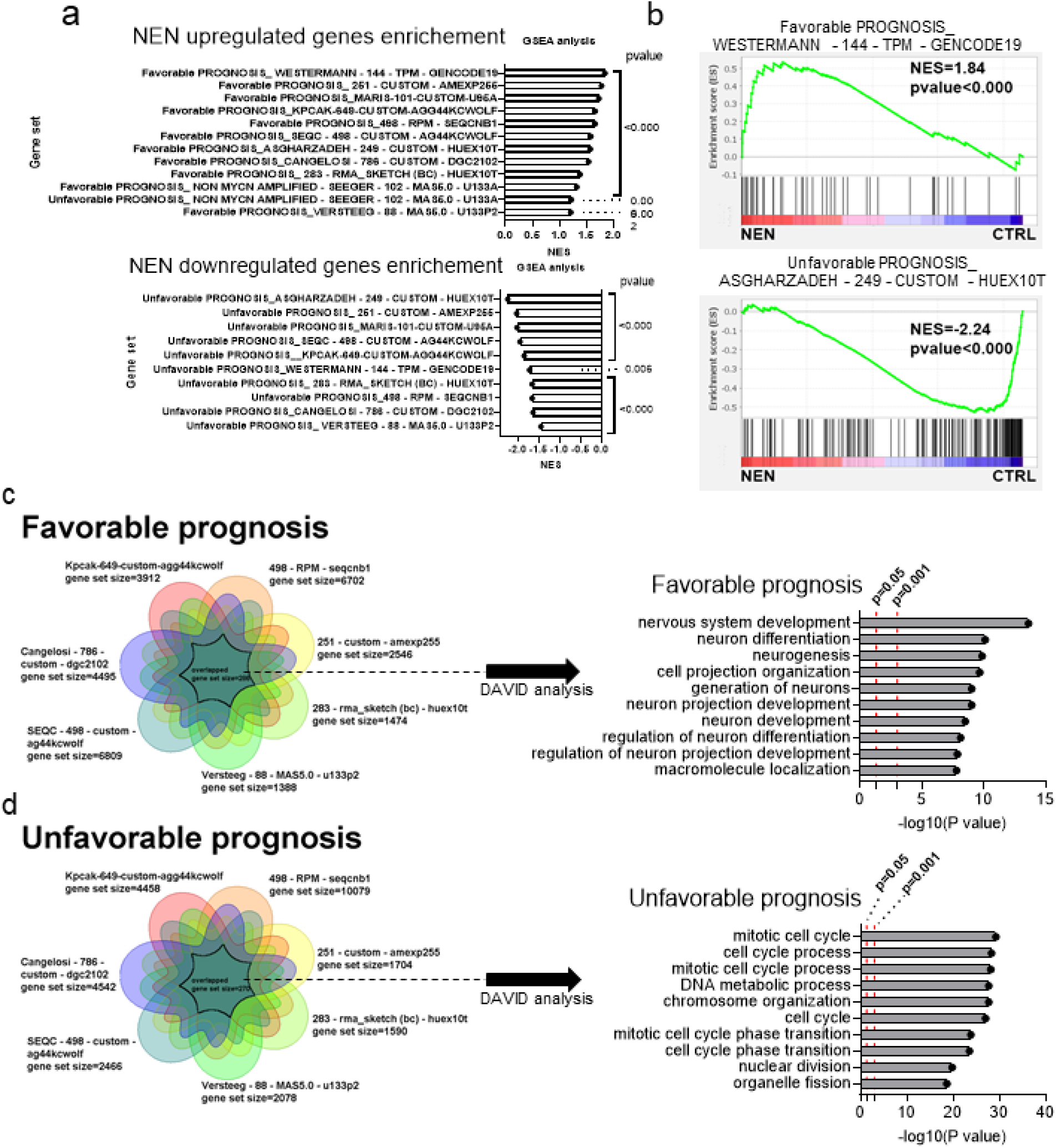
NEN showed favorable prognosis potential. (a) The gene expression data generated by RNA-seq(n=3) was analyzed by using GSEA to perform enrichment in SK-N-BE(2) cells. The gene sets (good or bad prognosis, pvalue<0.05) were defined from 11 available neuroblastoma databases from R2 (https://hgserver1.amc.nl/cgi-bin/r2/main.cgi). (b) Represented analysis plot from (a). (c) Overlap the good prognosis gene sets (p-value<0.05, gene number >1000) from 7 available neuroblastoma databases from R2 (https://hgserver1.amc.nl/cgi-bin/r2/main.cgi). And the overlapped genes were submitted to David analysis. (d) Overlap the bad prognosis gene sets (p-value<0.05, gene number >1000) from 7 available neuroblastoma databases from R2 (https://hgserver1.amc.nl/cgi-bin/r2/main.cgi). And the overlapped genes were submitted to David analysis.

## Discussion

Since Otto H. Warburg discovered that tumor cells have a high glucose consumption rate and produce large amounts of lactate in 1920s ^57^, oncologists have been interested in how tumor cells alters metabolic pathways to obtain advantages during cancer progression. In one of Warburg’s milestone reviews, he proposed that the cause of the Warburg effect was injury of respiration and as a consequence, cell dedifferentiation ^58^. At that time, although he was aware that the inhibition of respiration led to metabolic reprograming and insightfully proposed a connection between metabolic reprograming and cell dedifferentiation. However, the underlying mechanism was unclear due to the field’s limited understanding of metabolic control of epigenetics.

Inhibition of respiration is essentially the inhibition of the ETC, whose function is to oxidizes cellular NADH to NAD^+^. ETC inhibition leads to reduction of NAD^+^/NADH ratio. Low NAD^+^/NADH ratio not only drives lactate production from pyruvate to enhance the Warburg effect, but also disrupts TCA cycle flux, promoting reduction of α-KG to form L-2-HG ^29-30^. Similar to the D-2-HG produced from tumors carrying IDH1/2 mutation ^59-60^, L-2-HG also inhibits α-KG-dependent deoxygenases, including TET enzymes and JMJDs^61^, leading to DNA and histone hypermethylation^29, 62^. It was shown that this hypermethylation phenotype blocks cell differentiation, promoting tumor progression ^63-64^. Here we demonstrated that treatment using a mitochondrial uncoupler NEN effectively increased cellular α-KG levels in neuroblastoma cells to promote differentiation. NEN treatment increased NAD^+^/NADH ratio, which not only drives accelerated glutaminolysis to upregulate α-KG, but also blocks the conversion of α-KG to 2-HG. Together these data indicate that Warburg’s origin hypothesis was correct: the ETC activity is essential to cell differentiation. Surprisingly, even under hypoxia, NEN treatment is still effective in restoring α-KG and reducing 2-HG levels, suggesting that NEN supplementation has an advantage in demethylation *in vivo* over α-KG supplementation.

N-myc is the major oncogene amplified in NB. High N-myc expression is one key prognosis marker indicating poor survival in neuroblastoma patients. N-myc is downregulated upon induction of differentiation in neuroblastoma cells ^65^, suggesting a negative correlation between N-myc amplification and NB cell differentiation. Here we shown that mitochondrial uncoupler treatment could diminish the ‘undruggable’ N-Myc expression, highlighting the therapeutic potential in high grade neuroblastoma patients.

ETC inhibition not only promotes 2-HG production to reprogram the epigenome, but also induces reductive carboxylation flux. The citrate generated from α-KG provide acetyl-CoA for lipid synthesis, which is important for cell proliferation. Our data demonstrated not only that NEN treatment reversed this process, but it also increased cellular α-KG levels, suggesting that in addition to promoting proliferation by helping lipid synthesis, another important consequence of reductive carboxylation is to decrease intracellular α-KG levels and induce DNA hypermethylation.

In addition to αKG upregulation, mitochondrial uncoupler treatment caused global metabolic reprograming, including increasing NAD^+^/NADH ratio, AMP/ATP ratio, etc. These metabolic changes may further alter other signaling pathways that depends on NAD^+^ or ATP to remodel epigenetics and change cell fate. In summary, we have shown that NEN, a single agent, has extensive anti-tumor effects both *in vitro* and *in vivo*, by its ability to activate ETC. Upon ETC inhibition, altered metabolism causes global changes to epigenetic modification and signaling pathway rather than affecting a single gene or pathway. Thus, it is expected that activating ETC with mitochondrial uncouplers is an effective way of reversing this process and restoring the global metabolome and epigenome, which ultimately redirects the tumor cell into a differentiated state.

## Acknowledgement

This work was supported by a Stanford Maternal and Child Health Research Institute Research Scholar Award (2020) and an American Cancer Society Research Scholar Grant (RSG-20-036-01) to J.Y.

---

**Figure S1.**
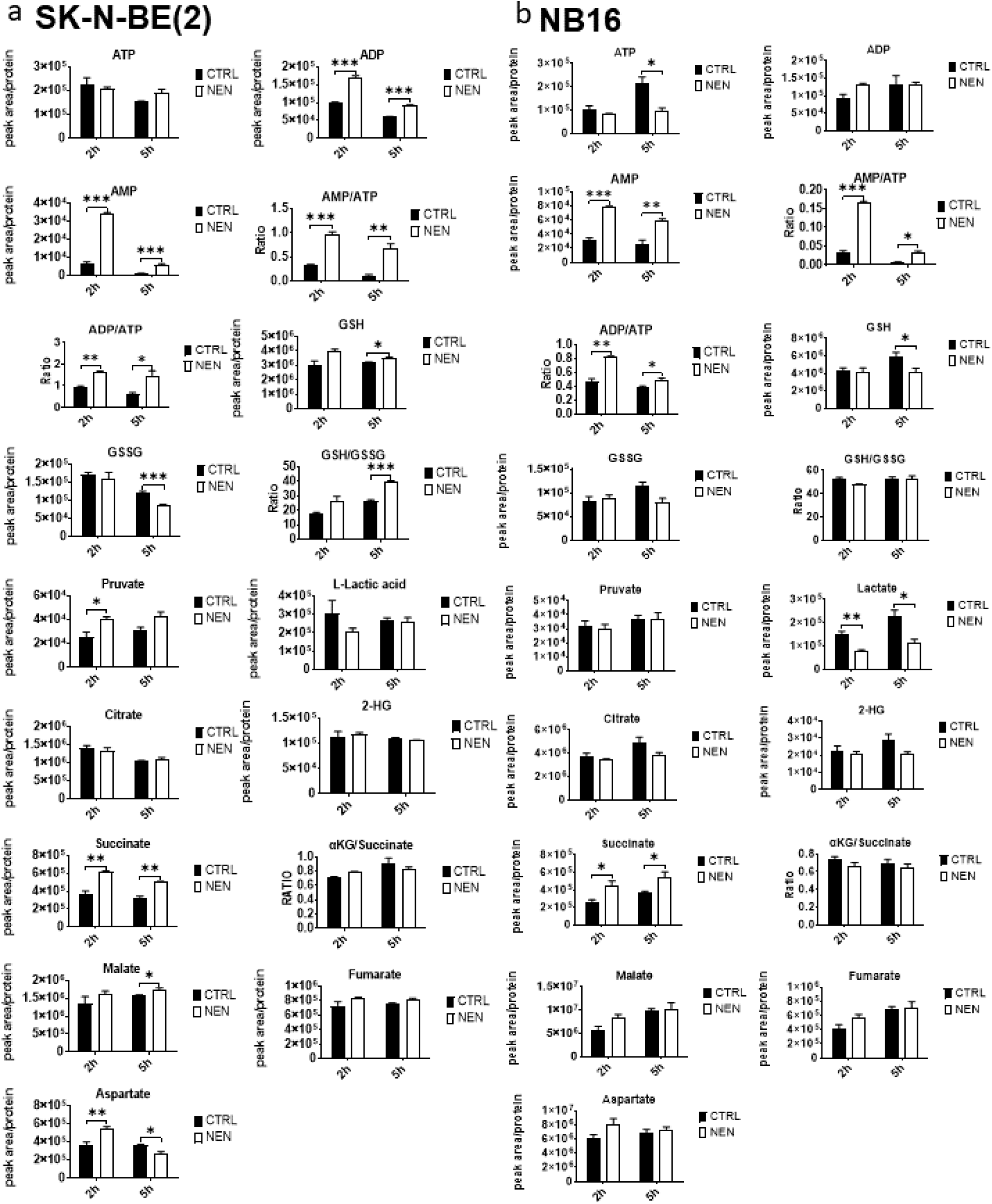
The metabolic profiling upon NEN treatment. Relative intracellular metabolite levels of the same samples in figure 1a and b. Data represent mean ± SEM (n = 3, biologically repeats). Representative of at least two independent experiments. *P < 0.05, **P < 0.01, ***P < 0.001. Two-sided Student’s t-test.

**Figure S2.**
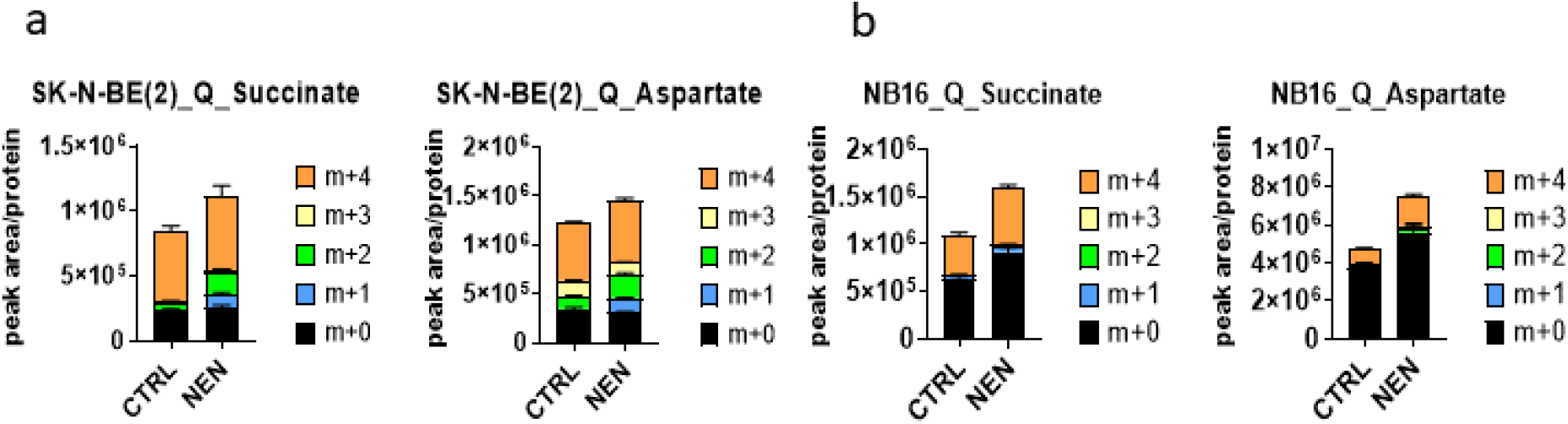
NEN treatment reprograms glutamine metabolism. (a) SK-N-BE(2) and NB16 cells were pretreated by DMSO or 1 μM NEN for 3h, then change the medium contained same treatment in the presence of U-1^3^C-glutamine for 2h. Data represent mean ± SEM (n = 3, biologically repeats). Representative of at least two independent experiments. *P < 0.05 **P < 0.01 and ***P < 0.001 Two-sided Student’s t-test.

**Figure S3.**
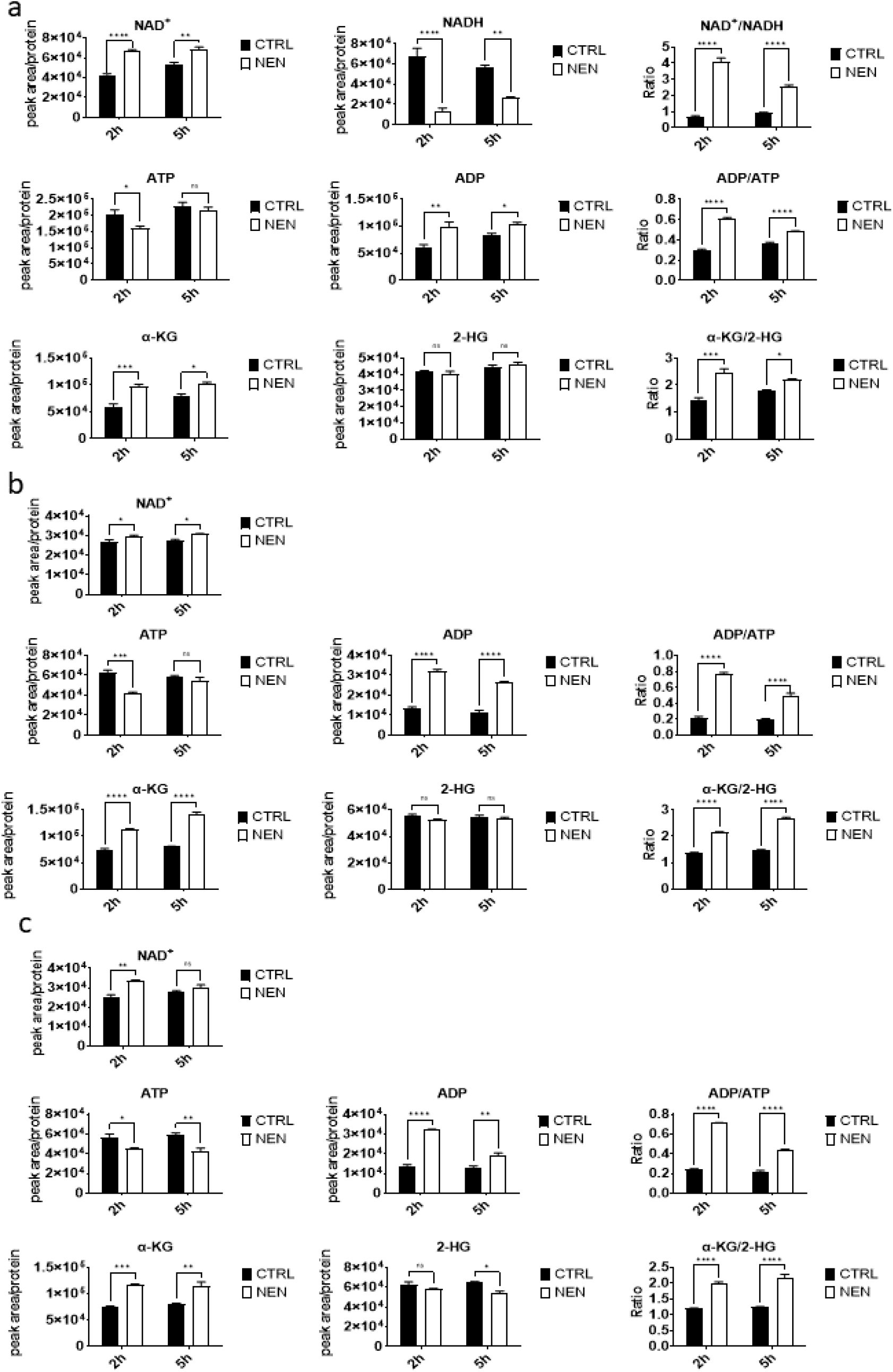
The metabolic reprograming effect of NEN on other cancer cell types. Relative intracellular metabolite level or ratios were measured using LC/MS in (**a**) Ovcar3 cells, (**b**)H29 and (**c**) H82 cells. Cells were treated with DMSO or 1 μM NEN for 2h or 5h. Data represent mean ± SEM (n = 3, biologically repeats). Representative of at least two independent experiments. *P < 0.05 **P < 0.01 and ***P < 0.001. Two-sided Student’s t-test.

**Figure S4.**
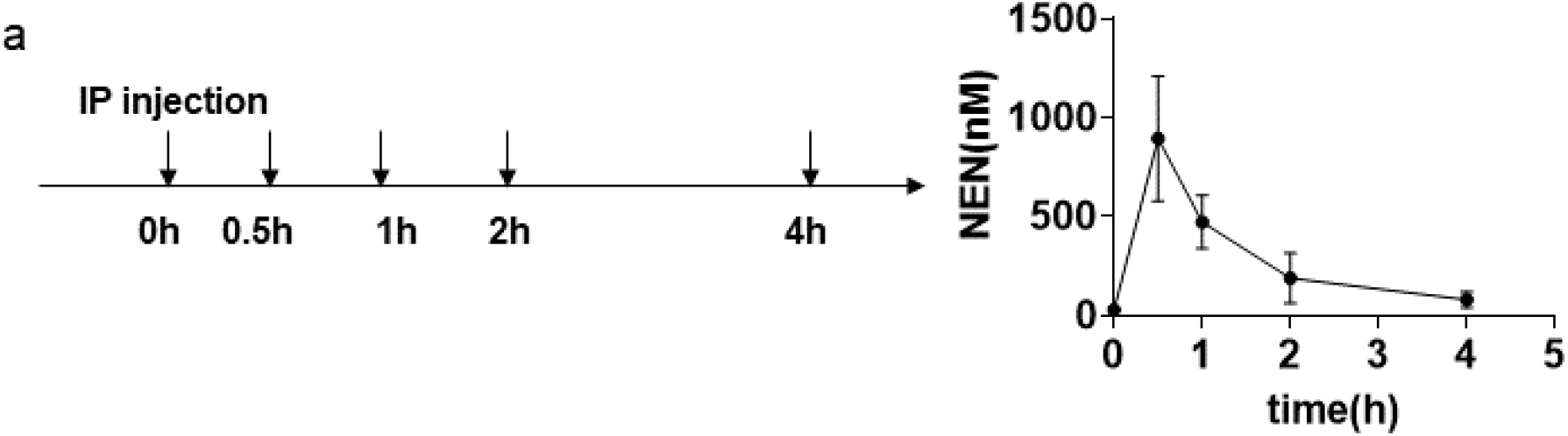
Plasma pharmacokinetics of NEN after through intraperitoneal injection. (a) 250μg NEN in DMSO were delivered to the mouses through intraperitoneal injection. Blood collection were performed for indicated time by tail vein sampling. The NEN concentration were measured by LC-MS (n=3).

